# The SARS-CoV-2 mutation landscape is shaped before replication starts

**DOI:** 10.1101/2022.09.29.510149

**Authors:** Diego Masone, Maria Soledad Alvarez, Luis Mariano Polo

## Abstract

Mutation landscapes and signatures have been thoroughly studied in SARS-CoV-2. Here, we analyse those patterns and link their changes to the viral replication niche. Surprisingly, those patterns look to be also modified after vaccination. Hence, by deductive reasoning we identify the steps of the coronavirus infection cycle in which those mutations initiate.

## Main

Modifications in the mutation landscape of a genomic sequence can result through several mechanisms^1^, such as error-prone polymerases, metabolism, and damaging agents, like an unbalanced redox environment. The comprehensive analysis of the SARS-CoV-2 interhost single base substitution (SBS) showed a mutational spectrum dominated by C>U and, surprisingly, G>U substitutions^2-5^. Our results confirmed that pattern (Fig.1A), followed by G>A and A>G, as expected. Transition-type SBS –the interchanges between purines (C>U and U>C) or pyrimidines (G>A and A>G)– were expected to be the most frequent, as they can result from the activity of antiviral enzymes such as APOBEC and ADAR deaminases in host cells^5-7^. Indeed, uracil is the outcome of cytidine deamination. In contrast, G>U transversions, particularly prevalent in SARS-CoV-2^8^, can result from stochastic processes, such as the misincorporation of nucleotides by an error-prone polymerase with a specific bias or the chemical modification of RNA. Those hypotheses have been discussed previously^3,4,9,10^, and it is widely agreed that G>U transversion is caused by mutagen exposure, like oxidation due to reactive oxygen species (ROS). That effect was observed not only in viruses but also in bacteria^11^. Thus, the process begins with the oxidation of a guanine base to produce 8-oxoguanine (8-oxoG). Like guanine, 8-oxoG can pair with cytosine; however, it can also pair with adenine (Fig.1B). Exceptionally, if 8-oxoG pairs with adenine during the first cycle of viral RNA replication, it can be substituted by uracil in the second replication cycle.

**Fig. 1.**
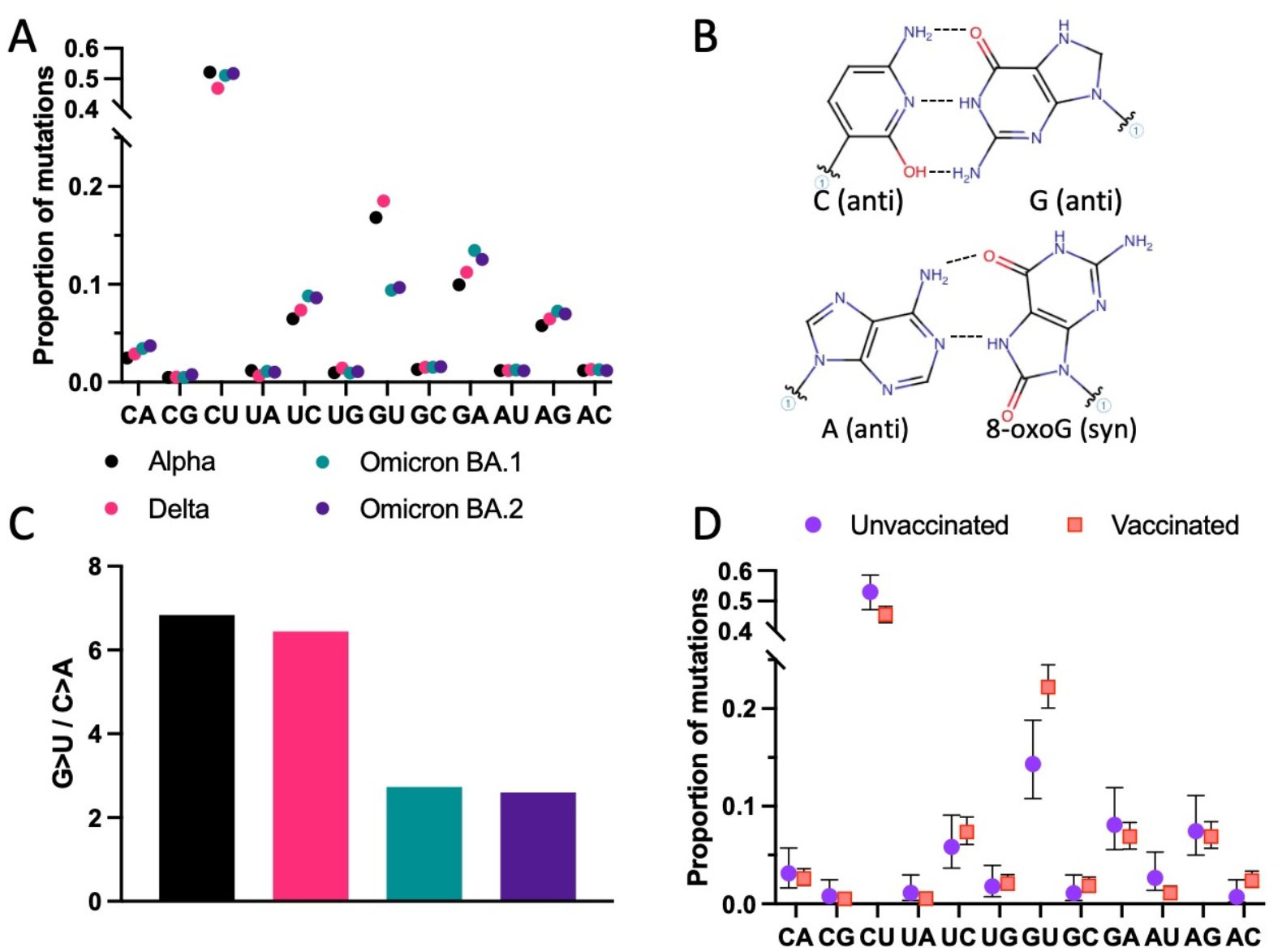
The environment of infected cells alters the G>U mutation incidence in SARS-CoV-2. (A) Analysis on mutation proportions across variants from ∼180,000 samples collected worldwide before 15th of January 2021, when less than 0.5% of the world population was vaccinated. The different colours denote the variants. (B) Diagrammatic representation of standard (Watson and Crick) pairing of guanine and cytosine (top panel) and Hoogsteen base pairing between 8-oxoguanine and adenine (bottom panel). (C) Proportion of G>U and C>A within the variants, coloured as in (A). (D) Comparison of single base substitution spectra of unvaccinated and vaccinated patients. Error bars denote confidence intervals (CI); however, in (A), CI are smaller than the size of the points.

Here, we considered how that misincorporation could occur during the intracellular life cycle of SARS-CoV-2. Various external mechanisms can explain modifications in the redox balance in infected cells, with the immune system as the prime suspect^12^. The immune cells and those regulating their functions vary through the respiratory tract^13,14^. Therefore, SBS patterns from those lineages infecting only part of the respiratory tract should differ from those that can infect the whole tract. For example, omicron subvariants (BA.1 and BA.2) mainly replicate in the upper respiratory tract (URT)^15^, which reflects in a significant decrease in the G>U/C>A ratio when compared to Alpha and Delta (Fig 1A and Fig1C; and Ruis *et al*.^16^). Furthermore, if the immune system were indeed responsible for the changes in the oxidative environment of the infected cells^12,17^, differences would be expected between SBS spectra from unvaccinated and vaccinated patients^18,19^. Therefore, we analysed samples from patients infected with alpha and delta variants divided into two groups, unvaccinated and vaccinated. Remarkably, there were significant alterations in the mutation landscape of SARS-CoV-2 genomes from vaccinated patients. Those alterations circumscribe to a rise in G>U transversion (Fig.1D), sustaining the importance of the immunological responses to the oxidative nature of those mutations.

Then, we hypothesised two scenarios where the nucleotide mispairing could occur when vRNA is outside or inside double-membrane vesicles (DMVs), leading to different substitution patterns (Fig.2A and 2B). SARS-CoV-2 contains a positive non-segmented RNA genome [(+)vRNA]. Its replication comprises the early translation of a large polypeptide, then cleaved to produce the RNA-dependent RNA polymerase (RdRp). Both (+)vRNA and RdRp are compartmentalised into DMVs^20^, avoiding the action of nucleases during vRNA replication^21^. vRNA is then processed through double-stranded RNA intermediates in a sophisticated manner involving (+)vRNA and (-)vRNA. Nevertheless, some vRNA molecules generated inside DMVs are transported to the cytoplasm to produce viral structural proteins. In this scenario, G>U and C>A should have similar magnitudes if the mispairing occurs during replication inside DMVs (Fig.2A). However, G>U substitutions prevail over C>A (Fig.1A, 1C and 1D), favouring the theory where the mispairing happens before the vRNA is enclosed into a DMV (Fig.2B). Subsequently, the asymmetry between G>U and C>A transversions can be explained by inferring that guanine oxidation occurs mainly outside DMVs (Fig.2A), so compartmentalisation can play a role in decreasing the exposure of vRNA to the oxidative environment, protecting it from ROS action. Similarly, other coronaviruses also shield their replication processes and machinery by using DMVs^22^. Consequently, it is unsurprising that the unbalance between G>U and C>A was previously observed in SARS-CoV and MERS-CoV^5^. Interestingly, the C>U/G>A ratio yields a value comparable to G>U/C>A (Fig.2C), suggesting that enzymatic deamination also happens before getting into the DMVs, and supporting the concept of the protective role of compartmentalisation.

**Fig. 2.**
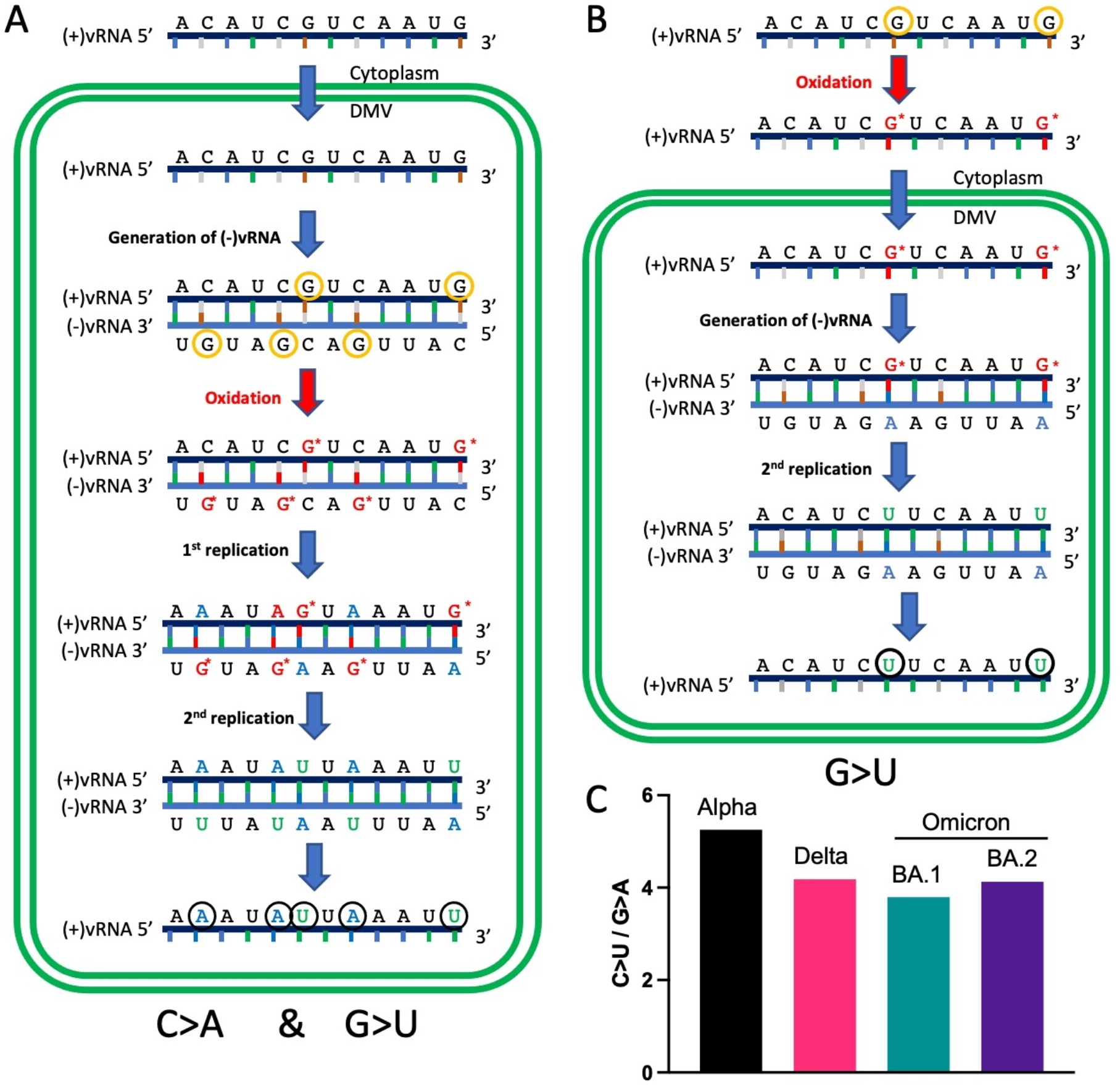
The influence of compartmentalisation on mutation patterns generated by oxidation. Differential pattern caused by mutagen exposure of vRNA guanines inside (A) or outside (B) double-membrane vesicles (DMVs), where vRNA replicates. Nucleotides circled in orange denote mutations that will occur in that scenario, while those in black mark the final product of the process. DMVs are delimited by double green lines. G* indicates 8-oxoguanine. (C) Proportion of C>U and G>A within the variants, coloured as in Fig.1A.

Additional studies are needed to elucidate in detail mechanisms driving viral mutation patterns and how that drive evolution of new SARS-CoV-2 strains. Particularly, if vaccines could cause novel strains appearance or to affect viral fitness through those mutations, further investigations are warranted to uncover how to manipulate that effect favouring their efficacy.

## Methods

### Calculation of SBS spectra

We calculated SBS mutational spectra for SARS-CoV-2 lineages using the 29th of July 2022 UShER SARS-CoV-2 phylogenetic tree^23^, as described in Ruis 2022^16^, with the small modifications described below.

To calculate SBS spectra for each vaccination status, we identified mutations on tip phylogenetic branches leading to sequences with known vaccination status in their GISAID metadata.

We calculated confidence intervals for mutation type proportions through the Wilson score interval using the total number of mutations as the number of trials and the proportion of the mutation type as the success proportion.

### Data availability

The findings of this study are based on metadata available on GISAID^24^, via gisaid.org/ EPI_SET_220927fu (doi: 10.55876/gis8.220927fu) and /EPI_SET_220925np (doi: 10.55876/gis8.220925np), for unvaccinated and vaccinated patients (respectively).

## Acknowledgements

The authors thank Dr Stuart Rulten, Prof Luis Mayorga and Prof Claudia Tomes for their critical and useful comments while conducting this study, and Dr Christopher Ruis (MRC-LMB, Cambridge, UK) and Dr Angie S. Hinrichs (University of California Santa Cruz, Santa Cruz, CA, USA), for their technical support. We gratefully acknowledge all data contributors, i.e., the Authors and their Originating laboratories responsible for obtaining the specimens, and their Submitting laboratories for generating the genetic sequence and metadata and sharing via the GISAID Initiative, on which this research is based. Funding: M.S.A. is a postdoctoral fellow of CONICET. This work was supported by grants from ANPCyT PICT2019-01889 and CONICET-PIP3195 to L.M.P.; ANPCyT PICT2020-01897 and CONICET-PIP0409 to D.M.

